# Kinome-wide activity classification of small molecules by deep learning

**DOI:** 10.1101/512459

**Authors:** Bryce K. Allen, Nagi G. Ayad, Stephan C. Schürer

## Abstract

Deep learning is a machine learning technique that attempts to model high-level abstractions in data by utilizing a graph composed of multiple processing layers that experience various linear and non-linear transformations. This technique has been shown to perform well for applications in drug discovery, utilizing structural features of small molecules to predict activity. However, the application of deep learning to discriminating features of kinase inhibitors has not been well explored. Small molecule kinase inhibitors are an important class of anti-cancer agents and have demonstrated impressive clinical efficacy in several different diseases. However, resistance is often observed mediated by adaptive Kinome reprogramming or subpopulation diversity. Therefore, polypharmacology and combination therapies offer potential therapeutic strategies for patients with resistant disease. Their development would benefit from more comprehensive and dense knowledge of small-molecule inhibition across the human Kinome. Because such data is not publicly available, we evaluated multiple machine learning methods to predict small molecule inhibition of 342 kinases using over 650K aggregated bioactivity annotations for over 300K small molecules curated from ChEMBL and the Kinase Knowledge Base (KKB). Our results demonstrated that multi-task deep neural networks outperform classical single-task methods, offering potential towards predicting activity profiles and filling gaps in the available data.

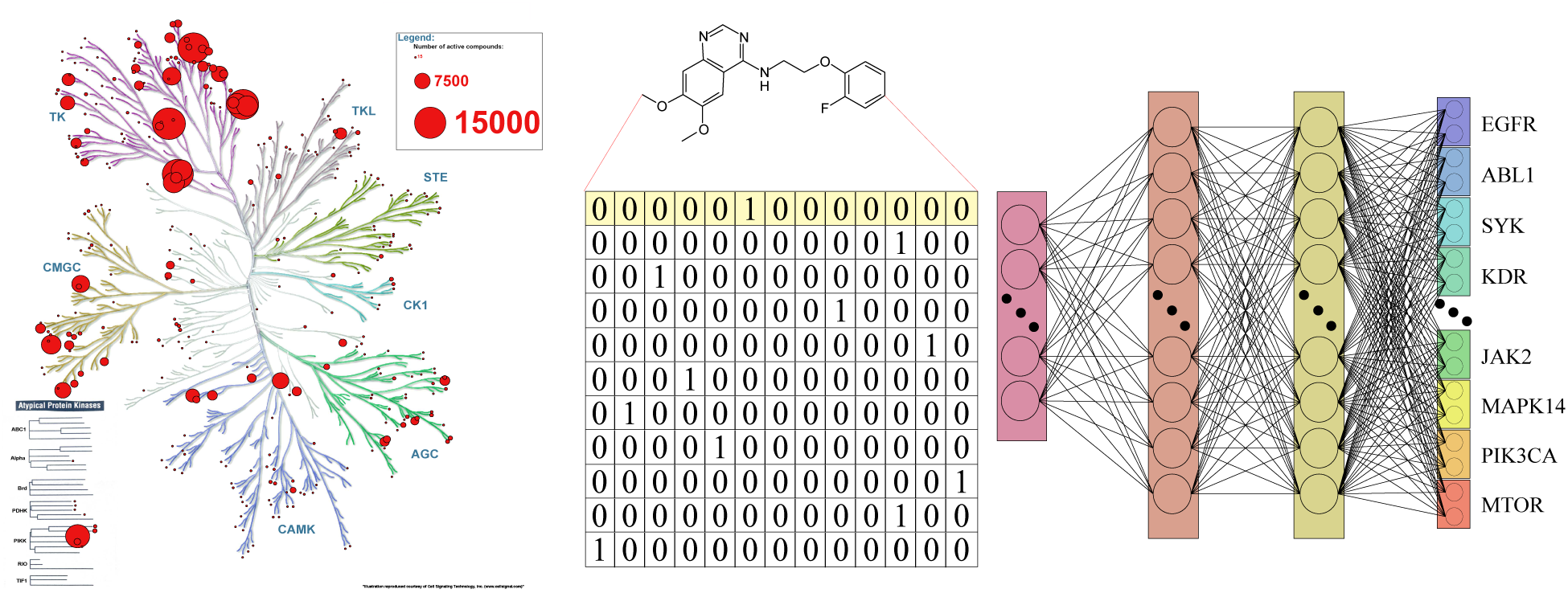
TOC Graphic

## INTRODUCTION

In the past decade, it has been increasingly well established that clinical benefit associated with drug treatment for a wide range of medical ailments varies substantially within patient populations^1–3^. This notion is especially palpable in the context of cancer therapy, where patient response to therapy is often poor, even with the advent of single-targeted therapies that address specific tumor genomic amplifications and disrupt oncogenic signaling processes. While patient initial response to targeted therapy is often observed, resistance is commonly detected as heterogeneity in the tumor cell population allows for adaptive selection of subpopulations that are not affected by the drug; or cells effectively rewire their signaling machinery to bypass the drug inhibition^4^. This paradigm has facilitated the acceptance of targeted polypharmacology and combination therapy strategies to address drug resistance and reduce recurrence in cancer patients.

Small-molecule kinase inhibitors are a main component of oncology drug development pipelines and are considered one of the most promising therapeutic strategies for cancer^5^. Biochemical and genetic studies have shown that many classes and types of cancer depend on protein kinase signaling mechanisms to facilitate cell proliferation, migration and survival. In addition, many kinases have shown to be targetable by small molecule drugs with over 30 having received FDA approval for treatment of specific cancers.^6^ However, response to these drugs has shown to be limited to a subset of patients, supporting the need for precision therapeutics approaches as we learn from the integration of clinical data, such as drug response and cancer multi-omics characterization, which features of these molecules are likely to give the best outcome^7–9^.

In-silico methods such as ligand-based virtual screening have been applied to kinase activity modeling and have shown that the utilization of known kinase small-molecule topological and bioactivity information can lead to the enrichment of novel kinase active compounds. The advantage of ligand-based virtual screening is that compounds can be readily evaluated for interactions across all kinases for which applicable amounts of bioactivity data are available. However, the method is not applicable if activity data are not available. If such models were sufficiently accurate, they could support the prioritization of compounds with a desirable polypharmacology profile while deprioritizing those compounds that may bind multiple therapeutically less relevant kinases or lead to toxicity in a specific disease or patient^10–11^.

Classification approaches to ligand-based virtual screening, also called target prediction or target fishing, involve a variety of single-task machine learning algorithms such as logistic regression, random forests, support-vector machines, and naïve Bayesian classifiers that aim to separate for each kinase target individually, whether a compound belongs in the active or inactive class^12–14^. Although these single-task methods perform relatively well in many instances, they do not take into consideration membership of molecules in multiple classes and therefore do not adaptively learn across different categories, limiting their applicability to predict profiles. To achieve better performance, machine learning methods must combine diverse sources of bioactivity data across multiple targets. This is especially relevant for kinases, where there is a large degree of similarity across many different kinases and their inhibitors.

Motivated by previous multi-task neural network architectures, we evaluated the performance of multi-task deep neural networks (MTDNN) on kinase classification as compared to single-task methods^15^. We compiled over 650K aggregated bioactivity annotations for over 300K unique small molecules spanning 342 kinase targets. We evaluated the machine learning methods using the reported actives and reported inactives and also using the reported actives and considering all other compounds in the global dataset as inactive. Our results indicate that multi-task deep learning results in substantially better predictive performance over single-task machine learning methods across three different 5-fold cross validation strategies. Additionally, the multi-task method continues to improve as more data are added, whereby single-task methods plateau or diminish in performance. Previous groups have applied deep learning to broad target classification and specific benchmarking datasets,^17^ however there has been no specific large-scale analysis for kinase targets.

## RESULTS

To investigate the implications and applicability of multi-task deep learning to kinase classification we implemented several experimental strategies to address these questions:

1. Can multi-task neural network architectures increase predictive performance of kinase classification compared to single-task methods?
2. Compared to single task methods, how well do multi-task models generalize (applicability domain) and how does their performance improve with more data or decrease with fewer data points?
3. Are multi-task neural network architectures applicable to domain-specific differences between kinase groups, i.e. do they perform across the Kinome despite large variations in available data?
4. Are the multi-task models applicable to new data, such as kinase profiling?

To address our first question, we built and evaluated single and multi-task classifiers using aggregated kinase bioactivity data obtained from ChEMBL^18^ and KKB^19^. This resulted in 668,920 activity data points distributed for 342 targets across the kinome and 317,228 unique compounds (Figure 1, Supporting Figure 1). First, known active – known inactive (KA-KI) models were evaluated. These models utilized only the reported active and inactive compounds for each task. Additionally, single and multi-task known active – presumed inactive (KA-PI) classifiers based on reported actives and considering all other compounds in the set as inactive, were built and evaluated to compare predictive performance across algorithms. Figure 2 shows the distribution of the area under the curve of the receiver operating characteristic (true positive rate over false positive rate) curve (ROC score) of each machine learning methods for all kinase tasks across different splitting strategies for MTDNN (see methods) and stratified random splitting for the single-task methods.

**Figure 1.**
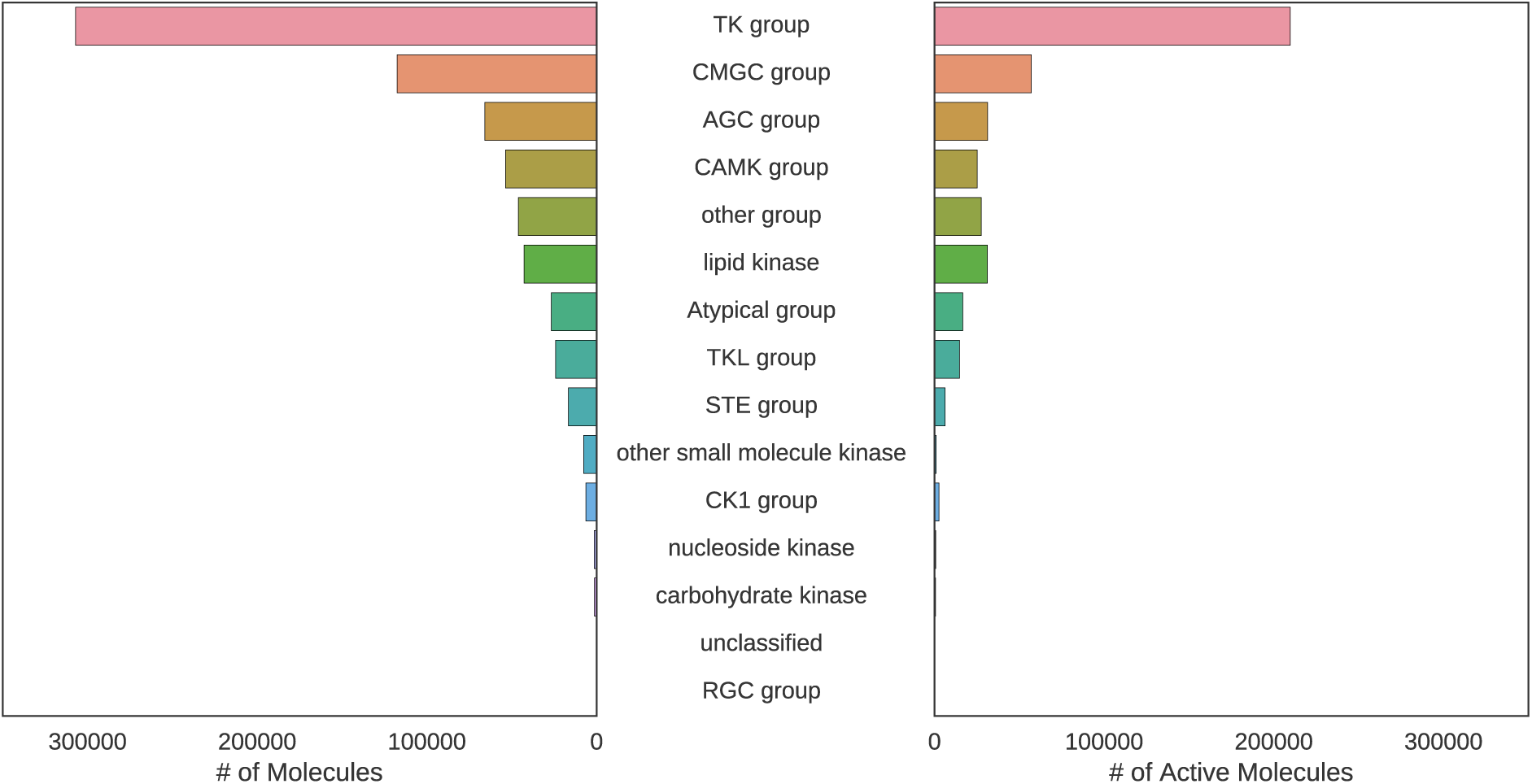
Distribution of dataset compounds by kinase family. The bar plot shows the sum of all molecules with kinase bioactivity annotations on the left and the sum of all active compounds for each kinase on the right shown for each kinase family.

**Figure 2.**
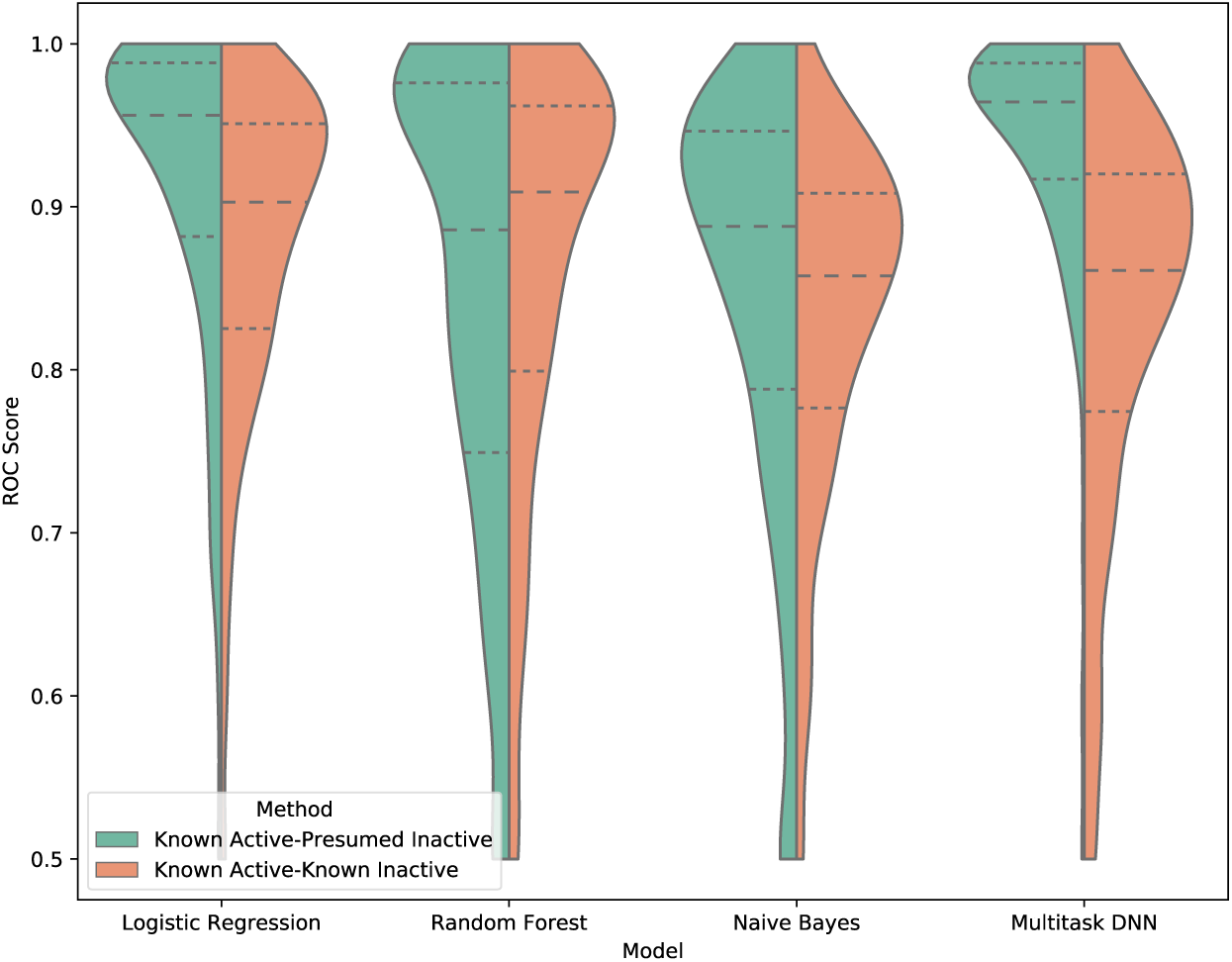
Differences in ROC score for kinase learning methods. Split violin plots with ROC score data distribution for all 342 kinase tasks. Each split violin plot represents a different modeling method with Known Active-Presumed Inactive (KA-PI) and Known Active-Known Inactive (KA-KI) datasets shown on the left and right, respectively. The receiver operating characteristic ROC area under the curve (AUC) is a metric to evaluate how well the model predicts true actives as actives vs the rate of true positives correlates with the rate of false positives. A higher score dictates a better discrimination of the model between true and false predictions.

Generally, the KA-PI models performed better than the KA-KI classifiers across all machine learning methods. Figures 2 and 3 clearly illustrates the best ROC scores were obtained by the KA-PI MTDNN model. Our best KA-PI multi-task model significantly outperformed single-task models across all cross-validation strategies and metrics. The MTDNN performed especially well for tasks containing large amounts of active compounds. These same trends were also observed when evaluating model performance based on enrichment factor (EF). However, it should be noted though that enrichment is of limited utility to characterize classifiers based on balanced datasets.

**Figure 3.**
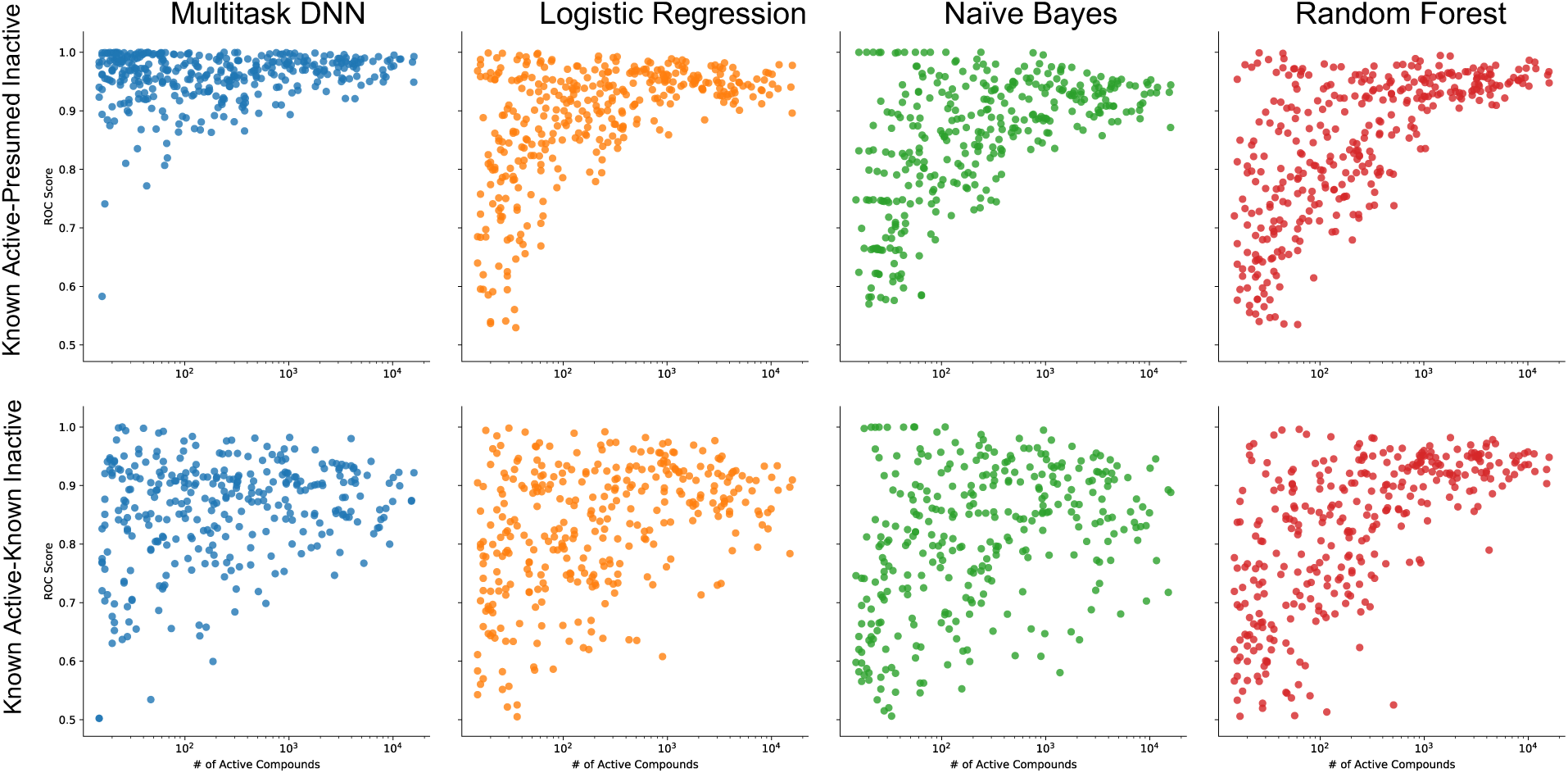
Effect of active compounds on kinase ROC score. Scatter plots ROC score data distribution for all 342 kinase tasks. Each box represents a specific model using either the Known Active-Known Inactive or Known Active-Presumed Inactive datasets.

In addition to improved performance with large datasets, compared to single-task methods, multi-task neural network predictions did not degrade as rapidly on tasks with smaller numbers of actives and continued to perform well even for tasks with <50 active compounds (Figure 3). Further, comparing the performance of models by enrichment factor at 0.1% of tested compounds across all kinase tasks, demonstrated that multi-task deep learning consistently performed above single-task classification methods for KA-PI (Figure 4). This is likely due to the utilization of relationships between targets and shared hidden representations among each kinase prediction task. As only the output weights are task specific, using data across all kinase targets can also help avoiding overfitting. Our hyperparameter optimization strategy demonstrated that for all kinase tasks, networks with two hidden layers of 2000 and 500 neurons resulted in the best performance across all metrics evaluated. Networks with three or more hidden layers did not provide any major improvements nor did they decrease in predictive performance. However, training networks with a larger batch-size did result in a decrease of performance. While a larger batch-size can reduce model training time, this increase in efficiency is not justified, because weighting parameters are not able to adequately tune themselves through each epoch.

**Figure 4.**
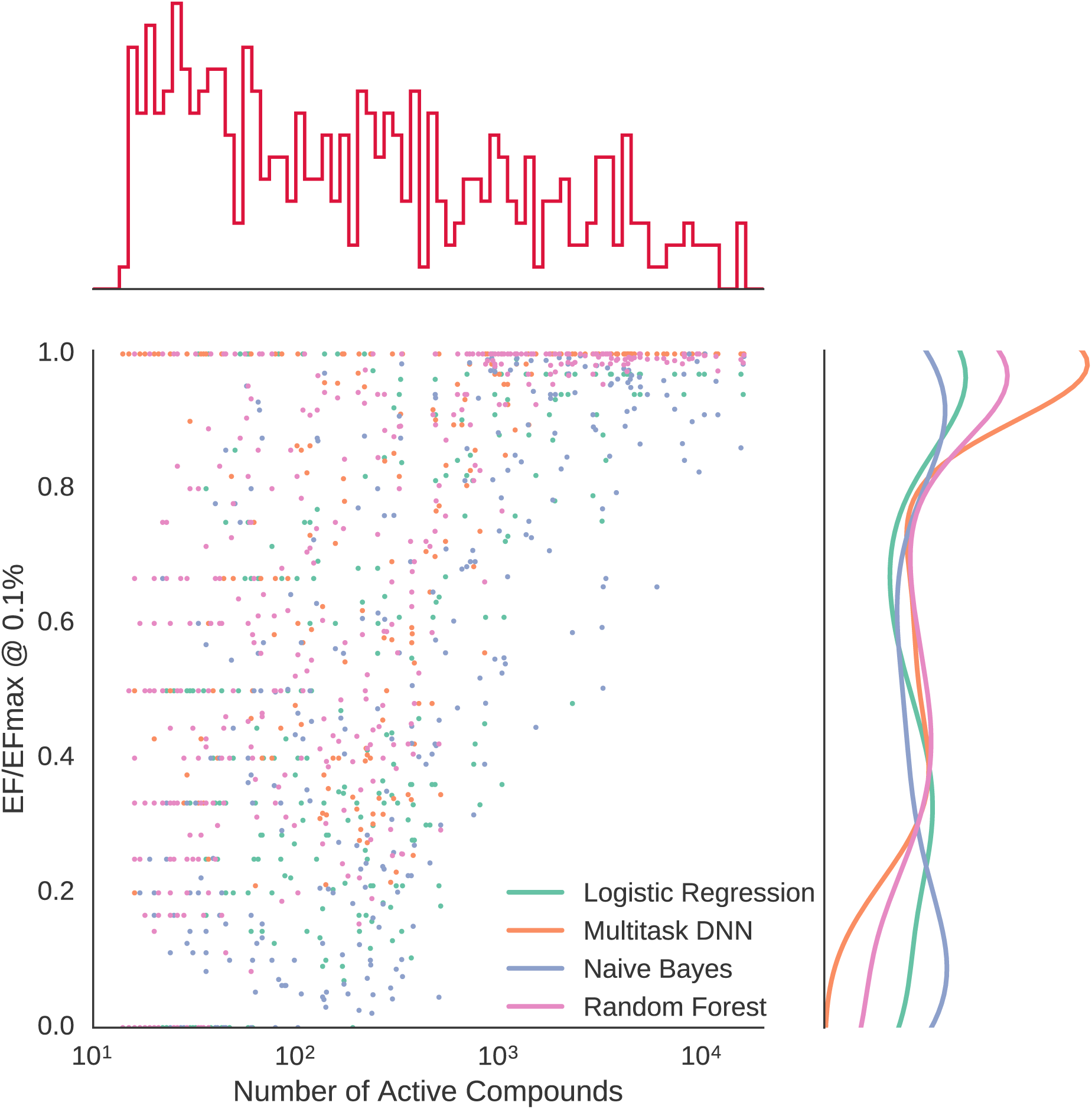
Normalized enrichment factor with respect to number of active compounds. Known active-presumed inactive (KA-PI) model performance across machine learning methods measured by normalized enrichment factor (EF/EFmax) as a function of number of active compounds for each kinase task.

Our goal was to develop predictors that are of practical utility for virtual screening or, in a perfect scenario for coarse virtual profiling. While it is ideal to build models only from confirmed actives and inactives (KA-KI approach), it is a recognized challenge that reported data in journal publications and patents, which underlie our datasets, are highly biased towards active compounds, reporting only few (often highly structurally similar) inactive compounds. In practice, as evident, for example, in experimental high-throughput screening (HTS) even of focused libraries, most compounds are inactive. Therefore, utilization of KA-KI classifiers – while highly predictive in many contexts – are of limited practical utility, especially in the context of multi-task learning methods. To overcome the limitation of not having enough known inactives, we considered all kinase compounds from ChEMBL and KKB that do not have reported activity for a given kinase as inactive for that kinase (KA-PI approach). While this may introduce some errors, they are likely not overwhelming, because it is common practice to report activity (including lack thereof) of the best compounds for the most similar kinases. In addition, profiling datasets are increasingly available and are part of ChEMBL and the KKB. To the extent to which presumed inactives introduce errors, we can assume that these will likely result in decreased model performance; therefore, the KA-PI approach appears reasonable to evaluate and compare machine learning methodologies and as more data become available and such errors are reduced, model performance should further increase.

Evaluation of the KA-PI models, which we considered more generally applicable, is shown in Figure 5 in which the average ROC scores of a five-fold cross-validation was calculated and binned by the number of active compounds. Comparing the performance of models of all kinase tasks demonstrated that multi-task deep learning consistently outperforms the single-task classification methods for KA-PI, but in particular performed much better for tasks with fewer active compounds. Indeed, the MTDNN continued to perform well even for tasks with less than 50 active compounds. Supporting Figure 2 illustrates that for kinase tasks with less than 500 active compounds, enrichment decreases for the single task modeling methods while the MTDNN is less affected. For classes with less than 50 active compounds, close to maximum enrichment (1000 for kinase tasks with less than 0.1% actives) is only obtained for the multi-task method.

**Figure 5.**
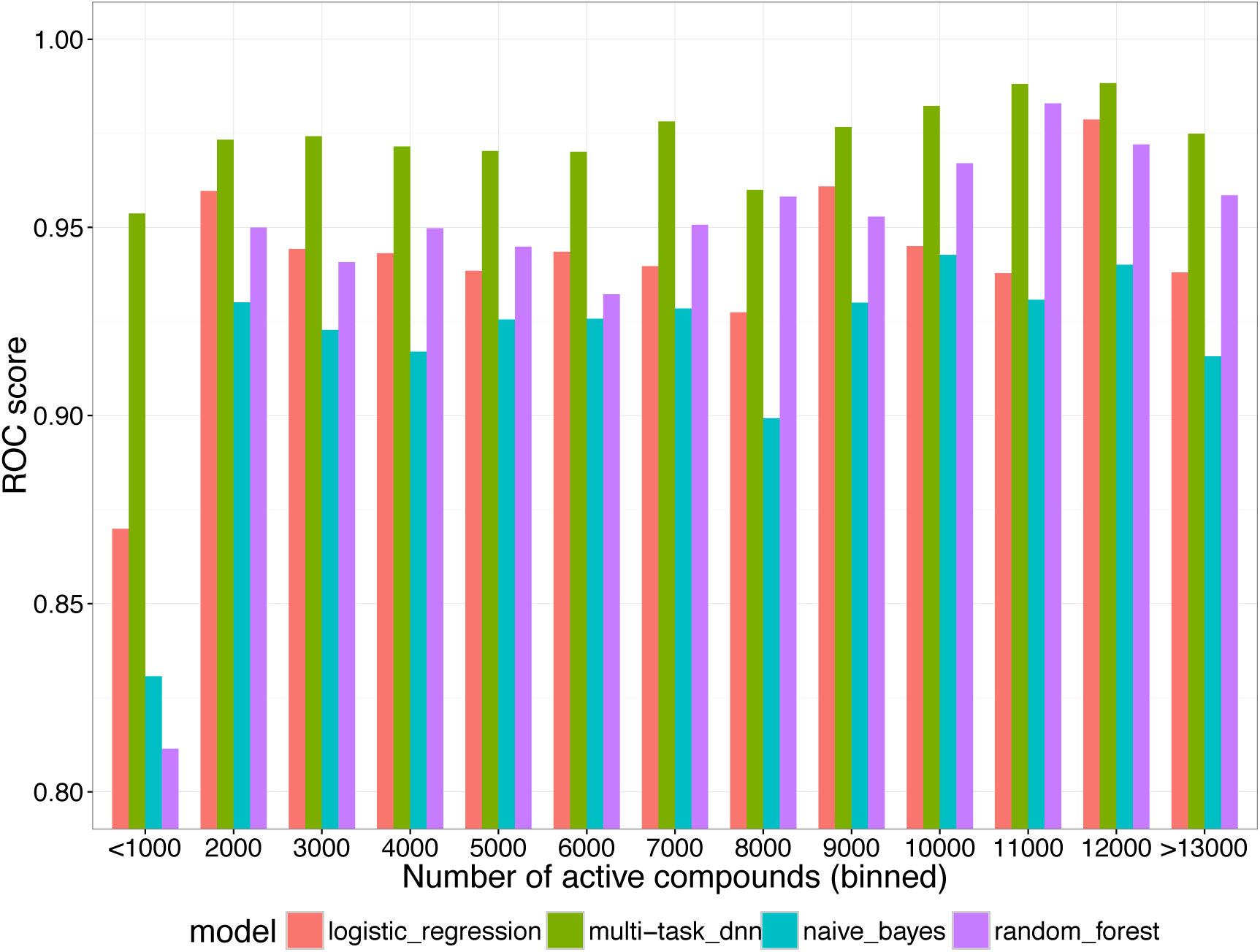
The effect of active compounds on ROC score across kinase learning methods. ROC score for different machine learning methods binned by ranges of active compounds across 342 kinase tasks (ROC score averaged over 5 repetitions of 5-fold random cross validation).

To further validate the KA-PI MTDNN, we randomized the kinase activity labels maintaining the number of actives for each dataset. 5-fold cross validation ROC scores around 0.5 were obtained. This corresponds to random classification, indicating the modeling approach did not overfit the data.

To further study how well the MTDNN models generalize, we evaluated the effect of different cross-validation strategies. Specifically, we implemented three different cross-validation strategies, including split based on chemical scaffold, by molecular weight, and randomly. We evaluated different strategies for splitting compounds by scaffold, including Murcko scaffold and clustering based on topological descriptors. We found that clustering our dataset into ~300 clusters with average ~1000 compounds each resulted in chemically diverse clusters (Supporting Figure 3); much better than, for example Murcko fragments. This was mainly due to the large number of Murcko scaffolds in our dataset (>100,000), which resulted in training and test sets that resembled random splitting. In each of the cross-validation strategies, the dataset was split into five subsets based on chemical scaffold / cluster, molecular weight and random order, respectively, and a model was trained five times, each time training on four sections, and evaluating on the held-out section. Hyperparameter optimization was utilized for all models and the evaluation and used half of the held-out section for validation and testing, respectively. As expected, ROC scores were much higher for models cross-validated by random splitting and worst for models evaluated by a different scaffold (Supporting Figure 4). As observed before, single tasks model performance decreased significantly for kinases with fewer active compounds, in contrast to the multi-task DNN approach, which continued to correctly classify active compounds even for those kinases that have a low number of actives. This trend was most pronounced with scaffold splitting, where many single-task methods failed to perform better than random when all instances of a specific scaffold were removed.

To illustrate the applicability of the MTDNN for virtual screening, the predictions for all active compounds over the different cross validation strategies were evaluated on how many true positives are recovered in 0.1% tested samples depending on the ratio of actives in a dataset (Figure 6). As expected, the fraction of true positives in a 0.1% test set on average increased with the ratio of active to total compounds and reached close to 100% as the latter reached above 0.1%. As the maximum possible enrichment is the inverse of the ratio of actives to total compounds, many of the models recovered close to all the actives in the test set. In practice that means, if 1000 compounds had been selected from one million, the best models would have identified between ten and close to 1000 actives. However, it should be noted that the best models assume the training set to be representative of the test set. Cross validation after splitting by scaffold or molecular weight performs much worse than random splitting. Nevertheless, even for new scaffolds, the models appear applicable for virtual screening.

**Figure 6.**
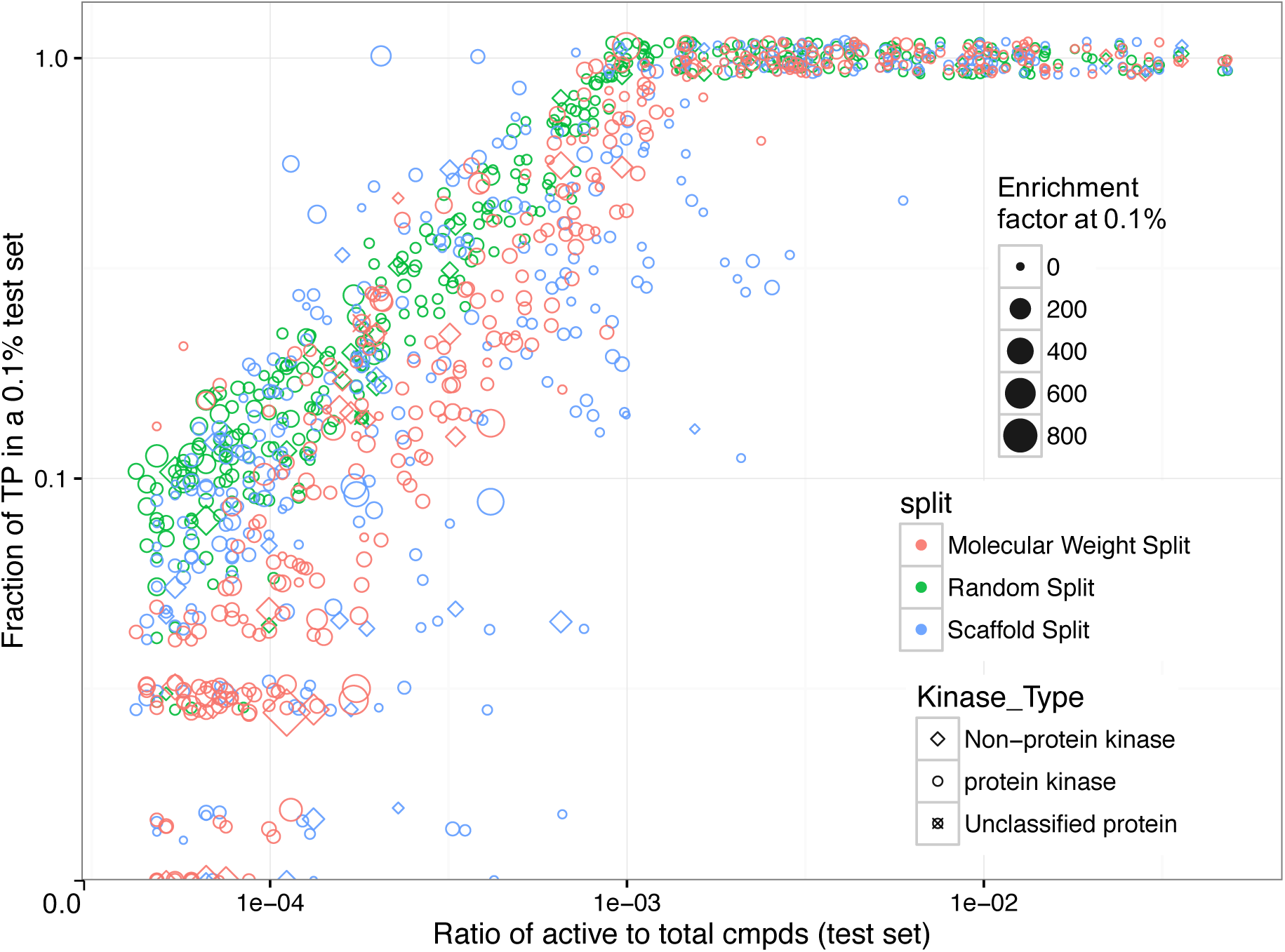
Influence of true positive at 0.1% to ratio of actives to total compounds in test set. Scatterplot with fraction of true positives at 0.1% of test set for all 342 KI-PI MTDNN kinase tasks across different cross validation splitting strategies shown as different colors. Each point represents a specific kinase task. Size indicates enrichment factor at 0.1%. Shapes correspond to non-protein, protein and unclassified kinases. Axes are log-log scale.

In addition to the different cross validation splitting strategies, we explored how model performance would scale with available data. To do that, for the top 100 kinase tasks with the most active compounds, we explored data dependence by randomly selecting 10, 20, 30 and 50% of the data. Evaluation of models across these different data thresholds revealed sustained increases in enrichment for the multi-task method, while single-task methods show much softer increases and eventually plateau or even start decreasing in performance (Supporting Figure 5).

To further characterize multi-tasks vs single task machine learning, we evaluated how similarities between datasets underlying the different kinase tasks affect model performance (Figure 7). All active compounds for each kinase task were compared globally to all other kinase task actives and the average maximum chemical similarities were calculated (see methods). Figure 7 shows the relationship of ROC score and chemical similarity to other kinase tasks. As Figure 7 illustrates, the multi-task DNN, in contrast to the single methods, performs particularly well for kinase tasks that share similar compounds with other kinase datasets. These results suggest that adaptively learning distributed representations of molecular features between active compounds for different classes that are very similar allows the model to better learn a distinct representation between all compounds, facilitating better predictive performance.

**Figure 7.**
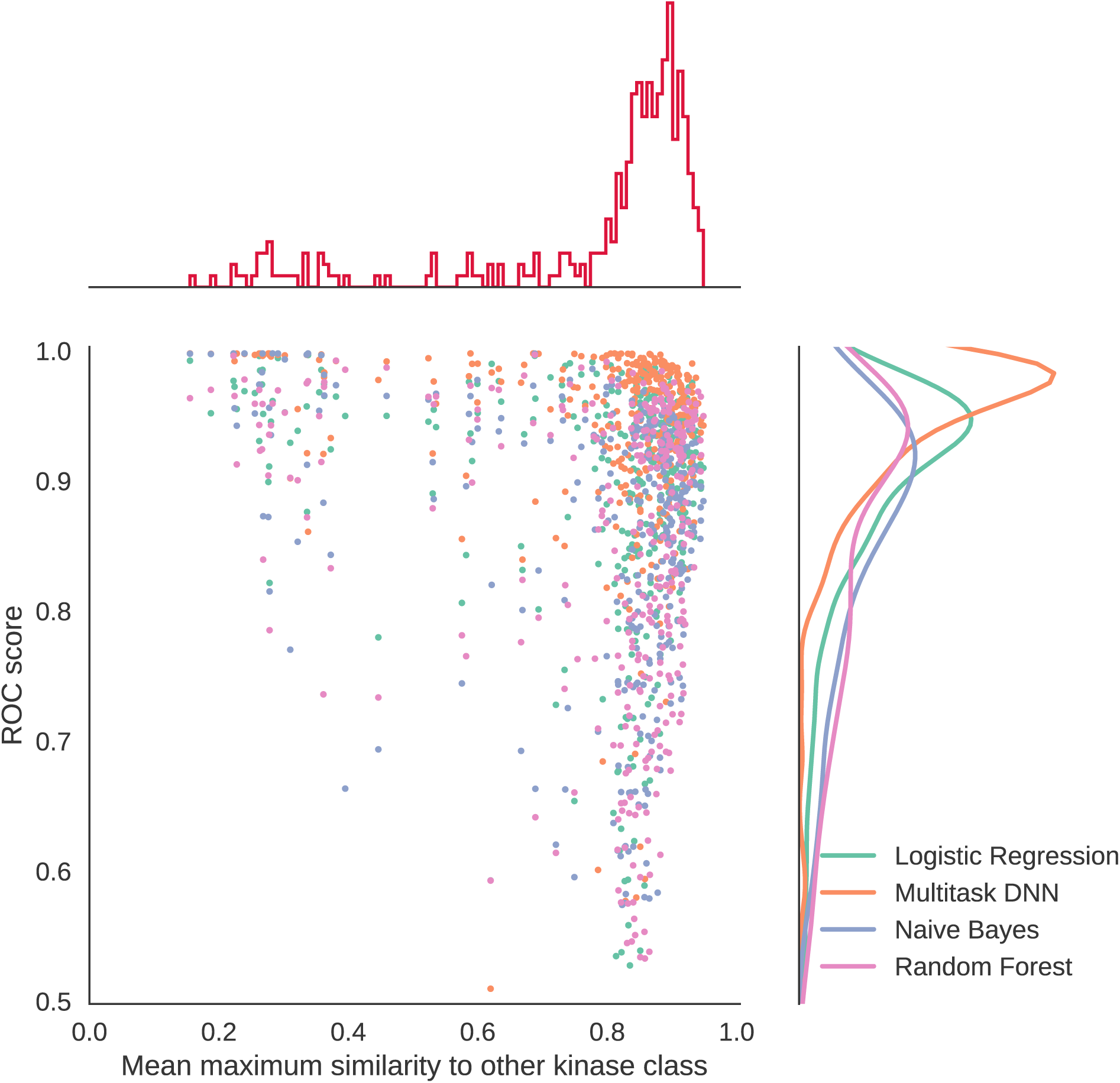
Effect of dataset similarities for multi-task vs single task methods. KA-PI model performance across machine learning methods measured by ROC score as a function of the average maximum similarity of compounds of one kinase tasks to all other kinase classes.

In the previous sections, we studied how the multi-task deep learning methodology performs compared to different single task machine learning methods with respect to the size and diversity of the datasets, how they generalize and their expected applicability to virtual screening. Another relevant aspect of Kinome-wide classification models is how they perform across the kinase target protein target family, as defined here by the kinase groups.^20^ Each kinase task was organized into its corresponding group (see methods) and model evaluation metrics were aggregated to discriminate differences in predictive performance. We observed that the multi-task models maintained high predictive performance across and within all kinase groups, even for those tasks that do not contain many active examples and are underrepresented globally (Supporting Figure 6). For single-task methods, there was a large amount of variance in the results across and within each kinase group tasks. As shown earlier, single-task methods performed well for kinase that had many active compounds, but poorly for those groups that did not. They performed on average better for those groups that have many kinase tasks with a large number of actives. However, the single tasks methods cannot leverage information from even the most similar kinases. In general, kinases within the same group are much more similar to one another, by sequence and small molecule activity, compared to kinases across different groups.^21^ These results thus support our earlier findings, that the MTDNN can cross leverage information from similar tasks; this appears to improve predictions for all tasks even for those that are underrepresented.

To further demonstrate how to apply our models to external datasets, we evaluated the performance of each KA-PI classifier at predicting compounds used in the LINCS KINOMEscan assay available from the LINCS Data Portal.^22^ Using purely chemical descriptors and overlap between our kinase tasks and kinases utilized in the assay, the multi-task deep learning model could correctly classify compound activity across kinases 90.8% of the time (Figure 7). The multi-task deep learning method was superior to other single task methods and was much better at predicting multiple activities across kinase tasks for compounds (see methods). This result is an important step forward in addressing sparsity in publicly available datasets. The results suggest the utility and broad applicability of our models, including virtual screening and progress towards virtual kinase profiling while overcoming some of the limitations resulting from sparse training data.

## METHODS

### Dataset Aggregation and Construction

We compiled our kinase target prediction dataset by obtaining all publicly available kinase bioactivity data from ChEMBL (release 21),^23^ a curated database of bioactivity measurements, and commercial kinase bioactivity data from KKB (release Q12016).^24^ Using kinase domain annotations from the Drug Target Ontology (DTO)^20^ we mapped domain information to UniProt, obtaining 485 unique UniProt identifiers. All UniProt IDs were mapped to ChEMBL release 21 target IDs and the corresponding KKB target ID. All bioactivity annotations obtained from ChEMBL were filtered by assay annotation confidence_score ≥ 5 and only compounds with activity annotations corresponding to a standard_type of Kd, Ki, IC50 or Activity were accepted. These data were then aggregated by chembl_id, standard_type, UniProt_id and the mean standard_value was calculated and −log_10_-transformed.

Commercial kinase bioactivity data from the Kinase Knowledge Base (KKB) was also obtained and aggregated by unique compound, endpoint, and target. ChEMBL and KKB data was then compared and overlapping compound and target annotations and endpoints were aggregated by their average. Compounds were grouped by canonical smiles and received an active label for each kinase where an aggregated pActivity value ≥ 6 was observed, and received no label otherwise. Kinase labels with < 15 unique active compounds were removed, leaving 342 unique kinase classes for which to build models.

ChEMBL and KKB often store chemical structures as published, therefore before combining and aggregating these datasets, we implemented an in-house chemical structure standardization protocol using Pipeline Pilot. Salts/addends and duplicate fragments were removed so that each structure consisted of only one fragment. Stereochemistry and charges were standardized, acids were pronated and bases deprotonated, and tautomers were canonicalized. For computing extended connectivity fingerprints of length 4 (ECFP4) descriptors for compounds (see below), stereochemistry and E/Z geometric configurations were removed (because these are not differentiated by standard ECFP fingerprints). This aggregation protocol yielded 668,920 measurements distributed across 342 targets and 317,228 unique compounds. Our datasets distribution was slightly skewed towards inactive compounds: 47% of the 668,920 training examples are active (about 320K). For further studies, we created two additional datasets, keeping the same number of total compounds but only keeping class labels for the 100 and 200 kinase classes with the most active molecules. The rationale behind gathering targets from public and commercial sources was to amass a collection of data that could leverage the power of both deep learning and multi-task algorithms. Given a set of related tasks, a multi-task network has the potential for higher accuracy as the learner detects latent patterns in the data across all tasks. This makes overfitting less likely and makes features accessible that may not be learnable by a single task.

We built the kinase predictors based on two different methods of handling kinase datasets described above. One method trained single and multi-task classifiers from known actives and known inactives (KA-KI). The other method built single and multi-task classifiers using all available data for unique kinase molecules, termed known active and presumed inactives (KA-PI). The models were built by identifying for each kinase task the active compounds and treating the remaining compounds as inactive (decoys). Both methods utilized all datasets and were characterized using the metrics introduced below.

### Small Molecule Topological Fingerprints (Features)

Extended connectivity fingerprints (ECFP4) were calculated using RDKit in Python. The ECFP4 algorithm assigns numeric identifiers to each atom, but not all numbers are reported. The fingerprints were hashed to a bit length of 1024, therefore very similar molecules can both be assigned the same numeric identifiers. Although increasing the number of bits reported can reduce the chances of a collision, there are also diminishing returns in the accuracy gains obtainable with longer fingerprints (e.g. 1024 bit, 2048 bit, or larger fingerprints can be used). This and computational complexity concerns were the pragmatic reasons why we chose to use 1024 bit ECFP4 fingerprints.

### Cross Validation Approach

To evaluate the predictive performance of the multi-task models, we implemented three different 5-fold cross validation strategies, including splitting by scaffold, molecular weight and randomly. In virtual screening, it is important to consider chemical diversity of the training and test sets for domain applicability and for evaluating how well the classifier generalize to new chemical space. For the scaffold-based cross validation, we performed hierarchical clustering for all compounds using Biovia Pipeline Pilot (version 16.1, 2016), specifying approximately 300 clusters with an average of approximately 1000 compounds. The pairwise Tanimoto similarities were calculated between all cluster centers and visualized to ensure that chemical dissimilarity was sufficient (Supporting Figure 1). Each cross validation held out 1/5 of the scaffolds and mean performance was calculated. Molecular weight was another distinguishing feature of compounds that could be used to estimate classifier performance. This method aims to keep the classifiers from overfitting on compound size and molecular weight can also be considered a simple surrogate for how different compounds are. Molecular weight was calculated in Python using RDKit. Compounds were sorted by increasing molecular weight and 1/5 of the dataset was held out each training iteration. Randomized 5-fold cross validation was also performed.

### Metrics and Model Evaluation

To adequately evaluate the machine learning models, we used a variety of metrics. The commonly used received operating characteristic (ROC) classification metric is defined as the true positive rate (TPR) as a function of the false positive rate (FPR). The TPR is the number of actives at a given rank position divided by the total number of actives, and the FPR is the number of inactives at a given rank position divided by the number of inactives for a given class. The area under the curve (AUC) is calculated from the ROC curve and is the metric we report. The calculation from a set of ranked molecules is given as follows:

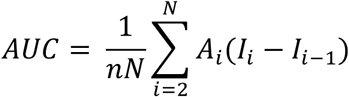

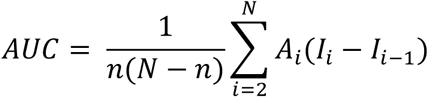

Where n is the number of active compounds, N is the total number of compounds, A is the cumulative count of actives at rank position i, and I the cumulative count of inactives at rank position i.

The enrichment factor (EF) is another metric we use and is very popular for use in evaluation of virtual screening performance. EF denotes the quotient of true actives among a subset of predicted actives and the overall fraction of actives and can be calculated as follows:

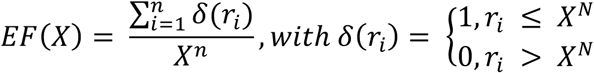

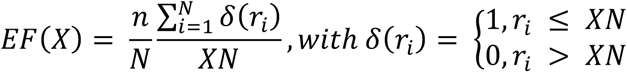

where r_i_ indicates the rank of the ith active, n is the number of actives, N the number of total compounds, X is the ratio within which EF is calculated. We evaluate the EF at 0.1% (X = 0.001) and 0.5% (X = 0.005) of the ranked test set. To evaluate the maximum achievable enrichment factor (EFmax), the maximum number of actives among the percentage of tested compounds is divided by the fraction of actives in the entire dataset. To more consistently quantify enrichment at 0.1% of all compounds tested, we report the ratio of EF/ EFmax. This normalized metric is useful because EF values are not directly comparable across different datasets due to the maximum possible EF being constrained by the ratio of total to active compounds and the evaluated fraction as shown in the equation above.

### Kinase Task Similarities

To evaluate how similar chemical structures in one kinase task are compared to those in all other kinase tasks, 25 diverse active compounds in each kinase task were selected using the diverse molecules component in Pipeline Pilot 9.0. This algorithm defines diverse molecules by maximum dissimilarity using the Tanimoto distance function and ECFP4 descriptors. For each kinase task, the average maximum Tanimoto similarity to all other kinase tasks was calculated based on the 25 diverse samples from each class. This task-based similarity is referred to the average maximum similarity of the reference class to all other kinase tasks as shown in Figure 7.

### Model Construction

Multi-task Artificial Neural Network Architecture

Neural networks can produce impressive non-linear models for classification, regression or dimensionality reduction and are applicable in both supervised in unsupervised learning situations. Neural networks take as input numerical vectors and render input to output vectors with repeated linear and non-linear transformations experienced repeatedly at simpler components called layers. Internal layers project distributed representations of input features that are useful for specific tasks.

More specifically, a multiple hidden layer neural network is a vector valued function of input vectors x, parameterized by weight matrices Wi and bias vectors bi,

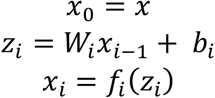

where fi are nonlinear activation functions such as rectified linear unit (ReLU) (max[0,zi]); xi is the activation of layer i and zi is the net input to layer i. After traversing L layers, the final layer x_L_ is output to a simple linear classifier, in our case the softmax, that provides the probability that the input vector x has label t:

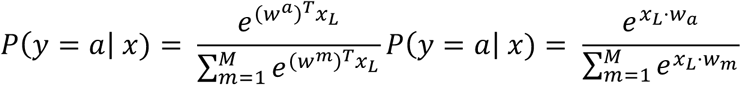

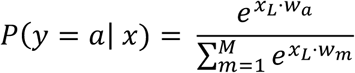

where M is the number of possible labels, in our case of binary prediction for each task (M = 2) and w_a_, w_m_ are weight vectors; x·w is the scalar (dot) product. Our network therefore takes a numerical chemical fingerprint descriptor of size 1024 as input, one or multiple layers of ReLU hidden units, and softmax output neurons for each kinase class or task. Given the known input and output of our training dataset, we optimized network parameters (x,y) = ({Wi,{bi) to minimize a defined cost function. For our classification problem, we used the cross-entropy error function for each task:

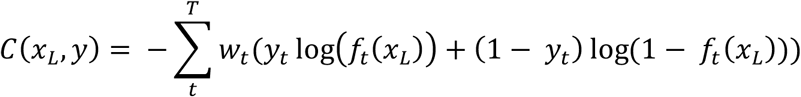

Where T is the total number of tasks, kinase classes in our implementation. The training objective was therefore the weighted sum of the cross-entropies over all kinase targets.

The algorithm was implemented in Python using the Keras package with Theano backend and was run on Nvidia Titan X GPUs with 12GB RAM to increase performance. Hyperparameter optimization included adjustments of momentum, batch size, learning rate, decay, number of hidden layers, number of hidden units, dropout rate, and optimization strategy. The best performing model consisted of training a batch size of 128 with two hidden layers of size 2000×500 using a dropout rate of 25% across each hidden layer for stochastic gradient descent learning. Model training varied in time from 2 days for all kinase classes to 4 hours for the 100 kinases with the most active compounds.

### Other single-task methods

All single-task methods were implemented in Python using the Sci-kit Learn machine learning library. Methods included logistic regression, random forests and Bernoulli naïve Bayes binary classifiers. Each method was implemented on the Pegasus supercomputer at the University of Miami (http://ccs.miami.edu/ac/service/pegasus/) using stratified 5-fold cross validation strategies and both KA-KI and KA-PI datasets, but training each class individually.

### KINOMEscan predictions

KINOMEscan datasets were obtained from the LINCS Data Portal (http://lincsportal.ccs.miami.edu/). The datasets contained 102 compounds with defined/known chemical structure and 486 different kinase targets (including mutations); however not all compounds were tested against all targets. Kinase activity was screened at 10 µM compound concentration. Kinase domains were curated and standardized using the Drug Target Ontology (DTO, http://drugtargetontology.org/) and datasets were joined into a data matrix of unique kinase domains and standardized small molecule canonical smiles; null values were introduced where no data was available. Kinase activity values were binarized for each small molecule kinase inhibition value [0,1] where 1 indicates active (≥50% inhibition) and 0 indicates inactive. KINOMEscan kinase domain targets were mapped to UniProt IDs via the DTO. UniProt kinase targets are mapped to ChEMBL and the KKB. To compare predicted kinase activity to the Kinomescan profiling results, we used the predicted probability to evaluate an active/inactive prediction.

## CONCLUSION

In this study, we investigated the utility of multi-task deep neural network classifiers to predict kinase activity of small molecule across the human Kinome. Although recently deep learning appears quite overhyped as the incarnation of artificial intelligence, we were interested how this method was applicable to reasonably large, target-family focused datasets of diverse small molecule kinase inhibitors and how it compares to more classical single task machine learning methodologies. A systematic study appeared justified to characterize the performance of deep neural networks, which are considerably more computationally expensive to train and also to use, i.e. run predictions. We investigated the applicability of heterogeneous datasets of small molecule kinase inhibitors to develop such models and studied in detail how various characteristics of the datasets influence model performance. The applicability and reusability of large public datasets is of considerable interest, because of the wide diversity of screening technologies and data processing pipelines and available metadata annotations.^25^ For example assay methods, detection technologies, assay kits, reagents, experimental conditions, normalization, curve fitting, etc, which are described in BioAssay Ontology,^26^ vary considerably among the tens of thousands of protocols and thousands of laboratories that generated these datasets. In many cases these details are not available in a standardized format and have to be manually curated from publications or patents. Using several single and multi-task machine learning methods, our results suggest that these datasets are highly useful to predict molecular activity.

We utilized over 650 thousand kinase-related activity annotations, which were processed and aggregated from public and commercial data sources, and ultimately determined 342 kinase classification tasks spanning the entire human Kinome. Using these data, we studied a variety of machine learning methods, including logistic regression, random forests, naïve Bayes and multi-task deep neural networks. To explore how the utility of the models including how they generalize with respect to chemical structure diversity and number of available data points, different cross validation strategies were utilized. Our results demonstrated reliable predictive performance across most kinase tasks if a sufficient number of active compounds or structure-activity data points were available. The best results across most kinases tasks were obtained using multi-task deep neural networks on known actives and a large background set of kinase-family focused presumed inactive compounds (KA-PI). Compared to single task methods, the MTDNN performed better for all situations investigated, including tasks with very few data points, tasks with many active compounds, and compounds that are dissimilar to the training data. The MTDNN performed well across all kinase groups, which considerably vary with respect to available data. One of the reasons of the superior performance appears to be their ability to adaptively learn molecular feature representations across different classes capturing latent features across tasks, which is not possible in single tasks methods. This characteristic of MTDNN could be particularly beneficial in the case of kinase-focused datasets, because of the pronounced similarities of the kinase APT-binding sites and the consequential and well-established considerable cross-reactivity of many small molecule kinase inhibitors.

Although the MTNDD approach gave the best results and appears most generalizable, it has considerably higher computational cost. However, with the latest generation of GPUs and as available datasets increase in size and diversity, deep neural networks will become increasingly relevant to fulfill the promise of virtual screening.

In summary, our results demonstrated high predictive performance across the human Kinome and suggest that multi-task deep neural network models trained on the corpus of diverse available small molecule kinase activity data are applicable for practical virtual screening and represent a step towards virtual kinase profiling.

## ANCILLARY INFORMATION

### Supporting Information

The following files are available free of charge.

xxxxxx.pdf - PDF document containing supporting figures (.pdf)

## AUTHOR INFORMATION

### Corresponding Author

Stephan C. Schürer

Department of Molecular and Cellular Pharmacology, Miller School of Medicine, University of Miami, Miami, FL, US.

### Author Contributions

BKA. developed code, performed computational modeling and data analysis; data curation and data integration; BKA and SCS performed data curation and integration; SCS devised the project; SCS and NA advised the project; BKA and SCS wrote the manuscript.

## ACKNOWLEDGMENTS

The authors would like to thank ChemAxon for providing the academic research license for their Cheminformatics software tools including JChem for Excel and the Marvin tools. The authors thank OpeneEye Scientific Software for their academic research licenses. We thank Eidogen-Sertanty for access to the KKB. SCS acknowledges computational resources of the CCS Drug Discovery program.

## REFERENCES

1. Ali, G.; Chella, A.; Lupi, C.; Proietti, A.; Niccoli, C.; Boldrini, L.; Davini, F.; Mussi, A.; Fontanini, G., Response to erlotinib in a patient with lung adenocarcinoma harbouring the EML4-ALK translocation: A case report. Oncol Lett 2015, 9(4), 1537–1540.

2. Ansari, J.; Fatima, A.; Chaudhri, S.; Bhatt, R. I.; Wallace, M.; James, N. D., Sorafenib induces therapeutic response in a patient with metastatic collecting duct carcinoma of kidney. Onkologie 2009, 32 (1-2), 44–6.

3. Baik, C. S.; Wu, D.; Smith, C.; Martins, R. G.; Pritchard, C. C., Durable Response to Tyrosine Kinase Inhibitor Therapy in a Lung Cancer Patient Harboring Epidermal Growth Factor Receptor Tandem Kinase Domain Duplication. J Thorac Oncol 2015, 10(10), e97–9.

4. Stuhlmiller, T. J.; Miller, S. M.; Zawistowski, J. S.; Nakamura, K.; Beltran, A. S.; Duncan, J. S.; Angus, S. P.; Collins, K. A.; Granger, D. A.; Reuther, R. A.; Graves, L. M.; Gomez, S. M.; Kuan, P. F.; Parker, J. S.; Chen, X.; Sciaky, N.; Carey, L. A.; Earp, H. S.; Jin, J.; Johnson, G. L., Inhibition of Lapatinib-Induced Kinome Reprogramming in ERBB2-Positive Breast Cancer by Targeting BET Family Bromodomains. Cell reports 2015, 11(3), 390–404.

5. Cohen, P., Protein kinases--the major drug targets of the twenty-first century? Nature reviews. Drug discovery 2002, 1(4), 309–15.

6. Wu, P.; Nielsen, T. E.; Clausen, M. H., Small-molecule kinase inhibitors: an analysis of FDA-approved drugs. Drug Discov Today 2016, 21(1), 5–10.

7. Barouch-Bentov, R.; Sauer, K., Mechanisms of drug resistance in kinases. Expert Opin Investig Drugs 2011, 20(2), 153–208.

8. Cecchin, E.; Agostini, M.; Pucciarelli, S.; De Paoli, A.; Canzonieri, V.; Sigon, R.; De Mattia, E.; Friso, M. L.; Biason, P.; Visentin, M.; Nitti, D.; Toffoli, G., Tumor response is predicted by patient genetic profile in rectal cancer patients treated with neo-adjuvant chemo-radiotherapy. Pharmacogenomics J 2011, 11(3), 214–26.

9. Zhou, L.; Shi, H.; Jiang, S.; Ruan, C.; Liu, H., Deep molecular response by IFN-alpha and dasatinib combination in a patient with T315I-mutated chronic myeloid leukemia. Pharmacogenomics 2016.

10. Schwab, R.; Petak, I.; Kollar, M.; Pinter, F.; Varkondi, E.; Kohanka, A.; Barti-Juhasz, H.; Schonleber, J.; Brauswetter, D.; Kopper, L.; Urban, L., Major partial response to crizotinib, a dual MET/ALK inhibitor, in a squamous cell lung (SCC) carcinoma patient with de novo c-MET amplification in the absence of ALK rearrangement. Lung Cancer 2014, 83(1), 109–11.

11. Allen, B. K.; Mehta, S.; Ember, S. W.; Schonbrunn, E.; Ayad, N.; Schurer, S. C., Large-Scale Computational Screening Identifies First in Class Multitarget Inhibitor of EGFR Kinase and BRD4. Sci Rep 2015, 5, 16924.

12. Chen, B.; Harrison, R. F.; Papadatos, G.; Willett, P.; Wood, D. J.; Lewell, X. Q.; Greenidge, P.; Stiefl, N., Evaluation of machine-learning methods for ligand-based virtual screening. J Comput Aided Mol Des 2007, 21 (1-3), 53–62.

13. Klon, A. E., Bayesian modeling in virtual high throughput screening. Comb Chem High Throughput Screen 2009, 12(5), 469–83.

14. Ma, X. H.; Jia, J.; Zhu, F.; Xue, Y.; Li, Z. R.; Chen, Y. Z., Comparative analysis of machine learning methods in ligand-based virtual screening of large compound libraries. Comb Chem High Throughput Screen 2009, 12(4), 344–57.

15. Ramsundar, B.; Kearnes, S.; Riley, P.; Webster, D.; Konerding, D.; Pande, V., Massively multitask networks for drug discovery. arXiv preprint arXiv:1502.02072 2015.

16. Keenan, A. B.; Jenkins, S. L.; Jagodnik, K. M.; Koplev, S.; He, E.; Torre, D.; Wang, Z.; Dohlman, A. B.; Silverstein, M. C.; Lachmann, A.; Kuleshov, M. V.; Ma’ayan, A.; Stathias, V.; Terryn, R.; Cooper, D.; Forlin, M.; Koleti, A.; Vidovic, D.; Chung, C.; Schurer, S. C.; Vasiliauskas, J.; Pilarczyk, M.; Shamsaei, B.; Fazel, M.; Ren, Y.; Niu, W.; Clark, N. A.; White, S.; Mahi, N.; Zhang, L.; Kouril, M.; Reichard, J. F.; Sivaganesan, S.; Medvedovic, M.; Meller, J.; Koch, R. J.; Birtwistle, M. R.; Iyengar, R.; Sobie, E. A.; Azeloglu, E. U.; Kaye, J.; Osterloh, J.; Haston, K.; Kalra, J.; Finkbiener, S.; Li, J.; Milani, P.; Adam, M.; Escalante-Chong, R.; Sachs, K.; Lenail, A.; Ramamoorthy, D.; Fraenkel, E.; Daigle, G.; Hussain, U.; Coye, A.; Rothstein, J.; Sareen, D.; Ornelas, L.; Banuelos, M.; Mandefro, B.; Ho, R.; Svendsen, C. N.; Lim, R. G.; Stocksdale, J.; Casale, M. S.; Thompson, T. G.; Wu, J.; Thompson, L. M.; Dardov, V.; Venkatraman, V.; Matlock, A.; Van Eyk, J. E.; Jaffe, J. D.; Papanastasiou, M.; Subramanian, A.; Golub, T. R.; Erickson, S. D.; Fallahi-Sichani, M.; Hafner, M.; Gray, N. S.; Lin, J. R.; Mills, C. E.; Muhlich, J. L.; Niepel, M.; Shamu, C. E.; Williams, E. H.; Wrobel, D.; Sorger, P. K.; Heiser, L. M.; Gray, J. W.; Korkola, J. E.; Mills, G. B.; LaBarge, M.; Feiler, H. S.; Dane, M. A.; Bucher, E.; Nederlof, M.; Sudar, D.; Gross, S.; Kilburn, D. F.; Smith, R.; Devlin, K.; Margolis, R.; Derr, L.; Lee, A.; Pillai, A., The Library of Integrated Network-Based Cellular Signatures NIH Program: System-Level Cataloging of Human Cells Response to Perturbations. Cell Syst 2017.

17. Unterthiner, T.; Mayr, A.; Klambauer, G.; Steijaert, M.; Wegner, J. K.; Ceulemans, H.; Hochreiter, S., Deep learning as an opportunity in virtual screening. Advances in Neural Information Processing Systems 2014, 27.

18. Bento, A. P.; Gaulton, A.; Hersey, A.; Bellis, L. J.; Chambers, J.; Davies, M.; Kruger, F. A.; Light, Y.; Mak, L.; McGlinchey, S.; Nowotka, M.; Papadatos, G.; Santos, R.; Overington, J. P., The ChEMBL bioactivity database: an update. Nucleic Acids Res 2014, 42 (Database issue), D1083–90.

19. Sharma, R.; Schurer, S. C.; Muskal, S. M., High quality, small molecule-activity datasets for kinase research. F1000Res 2016, 5.

20. Lin, Y.; Mehta, S.; Kucuk-McGinty, H.; Turner, J. P.; Vidovic, D.; Forlin, M.; Koleti, A.; Nguyen, D. T.; Jensen, L. J.; Guha, R.; Mathias, S. L.; Ursu, O.; Stathias, V.; Duan, J.; Nabizadeh, N.; Chung, C.; Mader, C.; Visser, U.; Yang, J. J.; Bologa, C. G.; Oprea, T. I.; Schurer, S. C., Drug target ontology to classify and integrate drug discovery data. J Biomed Semantics 2017, 8 (1), 50.

21. Davis, M. I.; Hunt, J. P.; Herrgard, S.; Ciceri, P.; Wodicka, L. M.; Pallares, G.; Hocker, M.; Treiber, D. K.; Zarrinkar, P. P., Comprehensive analysis of kinase inhibitor selectivity. Nat Biotechnol 2011, 29(11), 1046–51.

22. Koleti, A.; Terryn, R.; Stathias, V.; Chung, C.; Cooper, D. J.; Turner, J. P.; Vidovic, D.; Forlin, M.; Kelley, T. T.; D’Urso, A.; Allen, B. K.; Torre, D.; Jagodnik, K. M.; Wang, L.; Jenkins, S. L.; Mader, C.; Niu, W.; Fazel, M.; Mahi, N.; Pilarczyk, M.; Clark, N.; Shamsaei, B.; Meller, J.; Vasiliauskas, J.; Reichard, J.; Medvedovic, M.; Ma’ayan, A.; Pillai, A.; Schurer, S. C., Data Portal for the Library of Integrated Network-based Cellular Signatures (LINCS) program: integrated access to diverse large-scale cellular perturbation response data. Nucleic Acids Res 2017.

23. Papadatos, G.; Overington, J. P., The ChEMBL database: a taster for medicinal chemists. Future Med Chem 2014, 6(4), 361–4.

24. Schurer, S. C.; Muskal, S. M., Kinome-wide activity modeling from diverse public high-quality data sets. J Chem Inf Model 2013, 53(1), 27–38.

25. Stathias, V.; Koleti, A.; Vidovic, D.; Cooper, D. J.; Jagodnik, K. M.; Terryn, R.; Forlin, M.; Chung, C.; Torre, D.; Ayad, N.; Medvedovic, M.; Ma’ayan, A.; Pillai, A.; Schürer, S. C., Sustainable data and metadata management at the BD2K-LINCS Data Coordination and Integration Center. Nat Sci Data 2018, 5.

26. Abeyruwan, S.; Vempati, U. D.; Kucuk-McGinty, H.; Visser, U.; Koleti, A.; Mir, A.; Sakurai, K.; Chung, C.; Bittker, J. A.; Clemons, P. A.; Brudz, S.; Siripala, A.; Morales, A. J.; Romacker, M.; Twomey, D.; Bureeva, S.; Lemmon, V.; Schurer, S. C., Evolving BioAssay Ontology (BAO): modularization, integration and applications. J Biomed Semantics 2014, 5 (Suppl 1 Proceedings of the Bio-Ontologies Spec Interest G), S5.

